# Membrane association of VAMP2 SNARE motif in cells and its regulation by different lipid phases of synaptic vesicle membrane

**DOI:** 10.1101/587766

**Authors:** Chuchu Wang, Jia Tu, Shengnan Zhang, Bin Cai, Zhenying Liu, Shouqiao Hou, Zhijun Liu, Jiajie Diao, Zheng-Jiang Zhu, Cong Liu, Dan Li

## Abstract

Vesicle associated membrane protein 2 (VAMP2) contains a conserved SNARE motif that forms helix bundles with the homologous motifs of syntaxin-1 and SNAP25 to assemble into a SNARE complex for the exocytosis of synaptic vesicles (SV). Prior to SNARE assembly, the structure of VAMP2 is unclear. Here, using in-cell NMR spectroscopy, we described the dynamic membrane association of VAMP2 SNARE motif in mammalian cells at atomic resolution, and further tracked the intracellular structural changes of VAMP2 upon the lipid environmental changes. The underlying mechanistic basis was then investigated by solution NMR combined with mass-spectrometry-based lipidomic profiling. We analyzed the lipid compositions of lipid-raft and non-raft phases of SV membrane and revealed that VAMP2 configures distinctive conformations in different phases of SV membrane. The phase of cholesterol-rich lipid rafts could largely weaken the association of SNARE motif with SV membrane and thus, facilitate vesicle docking; While in the non-raft phase, the SNARE motif tends to hibernate on SV membrane with minor activity. Our work provides a spatial regulation of different lipid membrane phases to the structure of core SNARE proteins, which deepens our knowledge on the modulation of SNARE machinery.

## Introduction

Soluble N-ethylmaleimide-sensitive-factor attachment receptor (SNARE) complex is a macromolecular machinery which is largely involved in membrane fusion processes, in particular the fusion of synaptic vesicle (SV) membrane with pre-synaptic plasma membrane to release neurotransmitters^1^. The assembly of SNARE complex in neurons is driven by the formation of a stable four-helix bundle between VAMP2 (also known as synaptobrevin-2), syntaxin-1, and SNAP25, and modulated by multiple factors including proteins (e.g. synaptotagmin-1^2^, Munc18^3^, complexin^4^), lipids (e.g. sphingosine^5^, PIP2^6^) and metabolic ions (e.g. Calcium^7^). Among the core SNARE proteins, VAMP2 resides on the SV membrane, while the other two are on the plasma membrane. VAMP2 is composed of an extravesicular soluble domain and a C-terminal transmembrane domain (Fig. 1a). The soluble domain is intrinsically disordered and composed of an N-terminal proline-rich domain, a SNARE motif and a juxta-membrane domain. It is reported that the soluble domain of VAMP2 is unstructured in solution^8^and on lipid nanodisc^9^. While, in other lipid environments, the SNARE motif and juxta-membrane domain can associate with lipids, and this association varies largely from transient interaction to configuring an α-helix structure as proceeding from lipid bilayer to bicelle and micelle environments^10,11^. However, little is known about the structure and membrane association of VAMP2 extravesicular domains before vesicle docking in the native environment.

**Fig. 1.**
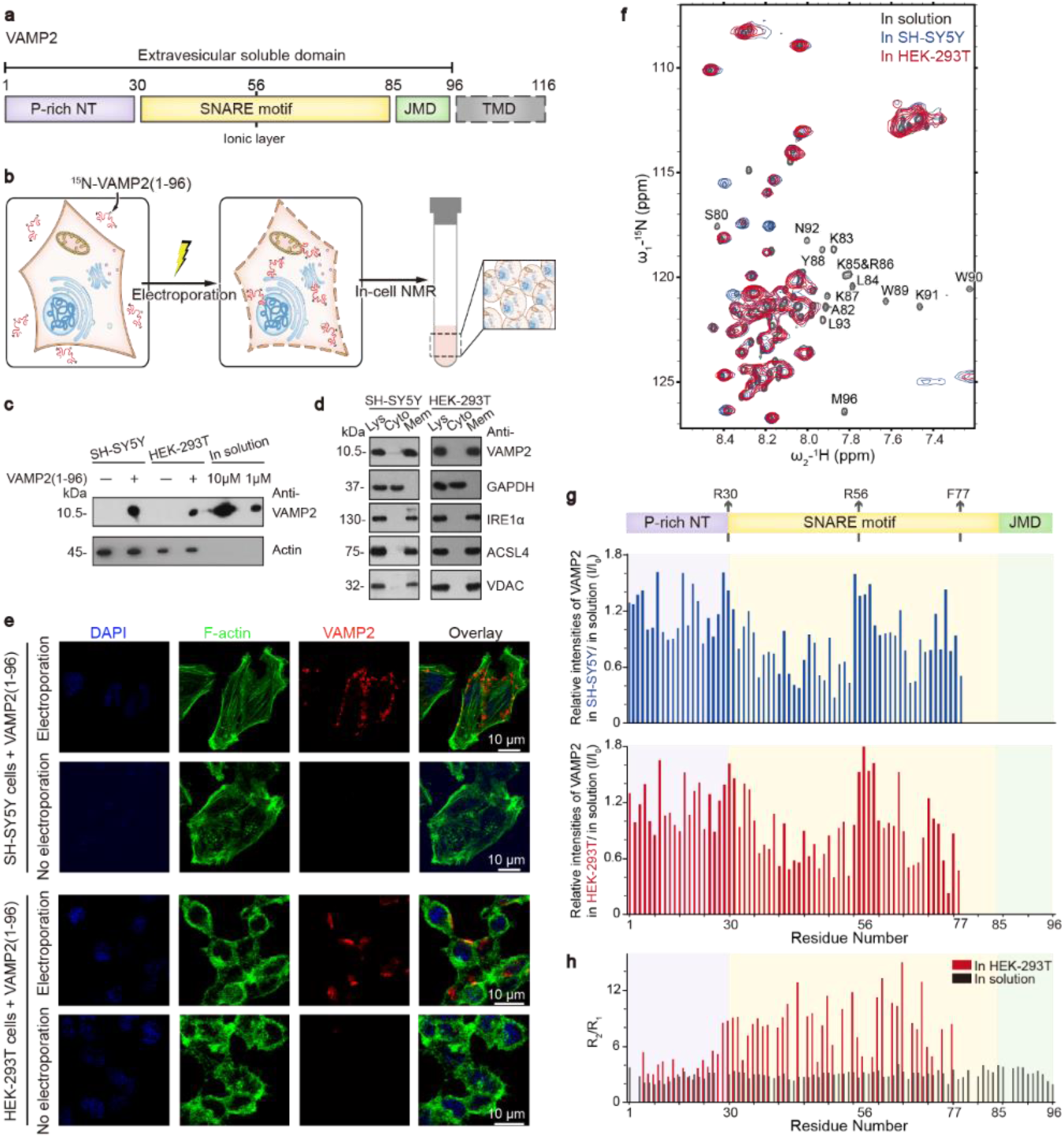
Membrane association of VAMP2 extravesicular domain in mammalian cells by in-cell NMR spectroscopy. **a**, Domain organization of VAMP2. P-rich NT: proline-rich N-terminal domain. JMD: juxta-membrane domain. TMD: transmembrane domain. R56 is conserved in the VAMP family which forms the zero ionic layer in the SNARE complex. **b**, Scheme of in-cell NMR sample preparation. NMR-visible ^15^N-VAMP2(1-96) was delivered into mammalian cells by electroporation. Un-delivered proteins were washed off before NMR signal acquiring. **c**, Estimation of VAMP2 quantity in cells by immunoblotting. Cells prepared for NMR were diluted 10 times before loading on gels. Known concentrations of VAMP2(1-96) proteins were loaded as standards. **d**, Sub-cellular localization of VAMP2(1-96) by fractionation and immunoblotting. Fractions of the total lysate (Lys), cytosol (Cyto) and membrane (Mem) were validated by immunoblotting with antibodies of GAPDH, IRE1α, ACSL4 and VDAC to indicate cytosol, endoplasmic reticulum membrane, plasma membrane-associated membrane and mitochondrial membrane, respectively. **e**, Sub-cellular localization of VAMP2(1-96) by immunofluorescence staining. F-actin filaments beneath cell membranes were stained by FITC-phalloidin. Nuclei were stained by DAPI. **f**, Overlay of 2D ^1^H-^15^N NMR spectra of VAMP2(1-96) in solution (black), SH-SY5Y cells (blue) and HEK-293T cells (red). Disappeared crosspeaks in cells were denoted. **g**, Residue-resolved relative NMR intensity ratios (I/I_0_) of VAMP2 (1-96) in cells to that in solution. Domain organization of VAMP2 extravesicular domain is indicated. **h**, Residue-resolved ratios of ^15^N transverse (R_2_, s^-1^) to longitudinal (R_1_, s^-1^) relaxation rates of VAMP2(1-96) in HEK-293T cells (red) and in solution (black).

In this study, we conduct in-cell NMR spectroscopy and observe that in the absence of transmembrane domain, VAMP2 remains associated with membranes in mammalian cells. In particular, the SNARE motif exhibits a dynamic interaction with native membranes, which can be adjusted by the changes of cellular cholesterol levels. We further dissect that the membrane association and activity of VAMP2 SNARE motif is elegantly tuned by the different lipid phases of SV membrane, which adds a spatial dimension on the complex modulation of SNARE machinery.

## RESULTS

### VAMP2 extravesicular domain associates with membranes in mammalian cells

To study the structure of VAMP2 in the native environment, we conducted in-cell NMR spectroscopy (Fig. 1b). We prepared ^15^N-labeled VAMP2(1-96) and electroporated it into HEK-293T cells and neuronal SH-SY5Y cells to cellular concentrations of ∼10 μM and ∼80 μM, respectively (Fig. 1c), which are comparable to the physiological concentration of VAMP2 (10∼100 μM) within synaptosomes of neurons^12^. The half-life of delivered VAMP2 in cells was about 18 hours (Supplementary Fig. 1). Interestingly, immunoblot showed that delivered VAMP2 is dominantly populated in the membrane fraction of the total cell lysates (Fig. 1d). Confocal microscopy further visualized that the delivered VAMP2 distributed along with the cell membrane skeleton in both cell types (Fig. 1e), which is in sharp comparison to that of α-synuclein identified to be evenly distributed in the cytosol in a previous in-cell NMR study^13^and in our experiment (Supplementary Fig. 2). These results demonstrate that, lacking of the transmembrane domain, VAMP2 can remain associated with membranes in cells.

**Fig. 2.**
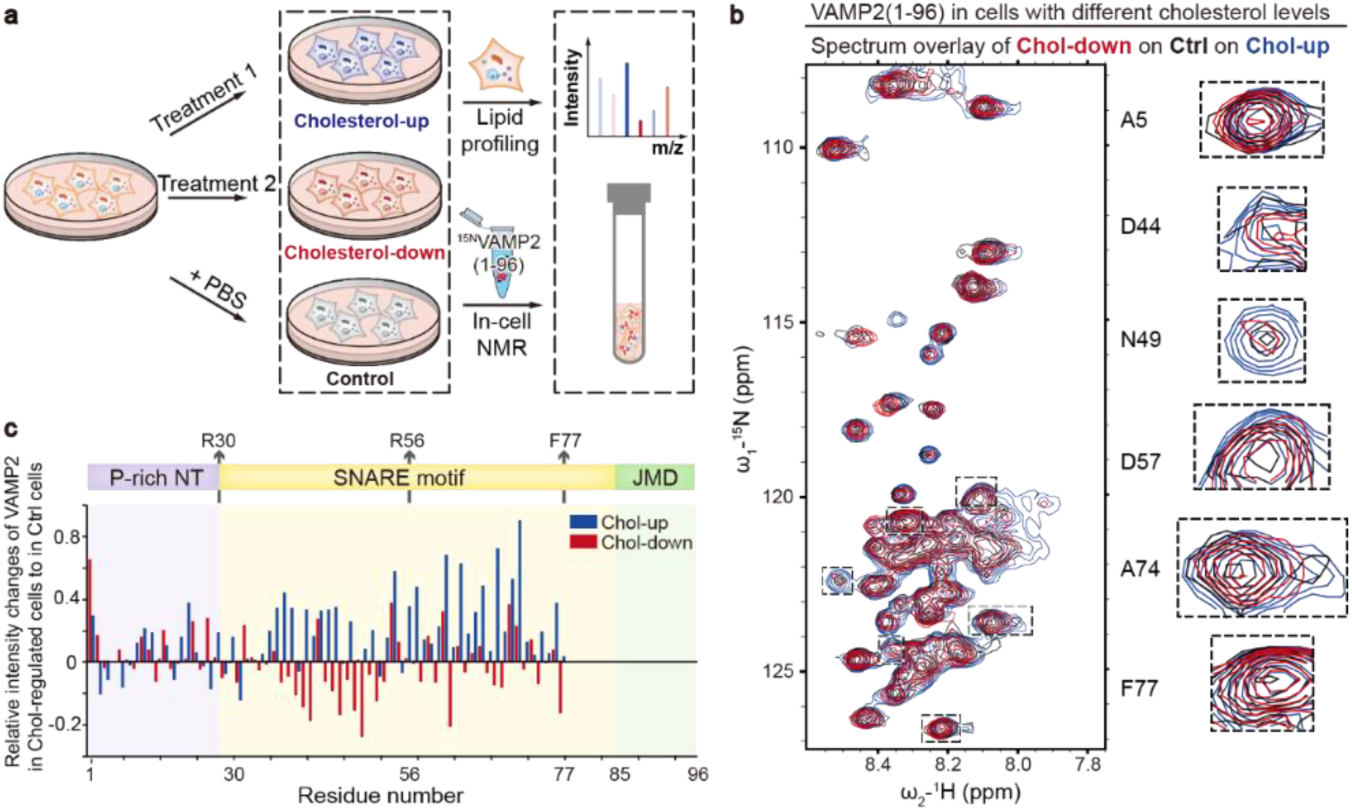
Residue-specific structural change of VAMP2(1-96) in cholesterol up-and down-regulated cells. **a**, Scheme of experimental design for studying the conformations of VAMP2 in cholesterol-level-regulated cells. Treatments 1 and 2 for increasing and decreasing cellular cholesterol levels were noted in the Supplementary Methods. **b**, Overlay of 2D ^1^H-^15^N NMR spectra of VAMP2(1-96) in cholesterol up-regulated (Chol-up, blue), down-regulated (Chol-down, red) and control (Ctrl, black) HEK-293T cells at the same contour levels. Representative crosspeaks were enlarged on the right. **c**, Relative NMR signal intensity changes of VAMP2(1-96) in Chol-up (blue) and Chol-down (red) cells to those in control cells. Details of data processing are described in the Supplementary Methods.

Next, we used SOFAST-HMQC pulse sequence to collect the signals of ^15^N-VAMP2(1-96) in cells. VAMP2 in HEK-293T cells exhibited a similar spectrum in terms of chemical shift and intensity, to that in SH-SY5Y cells (Fig. 1f, g), indicating that VAMP2 adopts a conserved conformation in these two different cell types. The residue-specific NMR crosspeaks of VAMP2 in cells, in comparison with those in solution, exhibited non-uniformly signal broadening and without obvious chemical shift perturbations, indicating that different regions of VAMP2 might interact divergently with cellular partners. The signal changes of VAMP2 in cells were not come from the crowding intracellular environment since crowding agents caused general signal attenuations of VAMP2, rather than regional signal changes (Supplementary Fig. 3). The 2D ^1^H-^15^N NMR spectra showed that in the intracellular environment, the signals of residues 78-96, which covers the juxta-membrane domain and a short C-terminal region of VAMP2 SNARE motif, completely disappeared (Fig. 1f, g). Deletion of this region (VAMP2(1-78)) resulted in a weakened membrane association with ∼25% of total VAMP2 released into cytosol (Supplementary Fig. 4), suggesting that the NMR signal missing of residues 78-96 in cells may result from membrane binding. NMR signals of the major region of the SNARE motif also exhibited significant signal attenuations (Fig. 1f, g). ^15^N transverse to longitudinal relaxation rates (R_2_/R_1_) further showed that residues 35-78 of the SNARE motif is less flexible and more ordered in cells than in solution (Fig. 1h). Truncation of the C-terminal half of the SNARE motif together with the juxta-membrane domain (VAMP2(1-59)), nearly completely abolished the membrane association of VAMP2 (Supplementary Fig. 4), which indicates that besides the juxta-membrane domain, the SNARE motif is also critical for the membrane association of VAMP2. Note that the conserved residue R56, which forms a “zero ionic layer” in the assembly of *trans*-to *cis*-SNARE complex^14,15^, exhibited no large signal changes (Fig. 1f, g). The N-terminal proline-rich domain also showed no significant NMR signal change in cells and in solution (Fig. 1f, g).

**Fig. 3.**
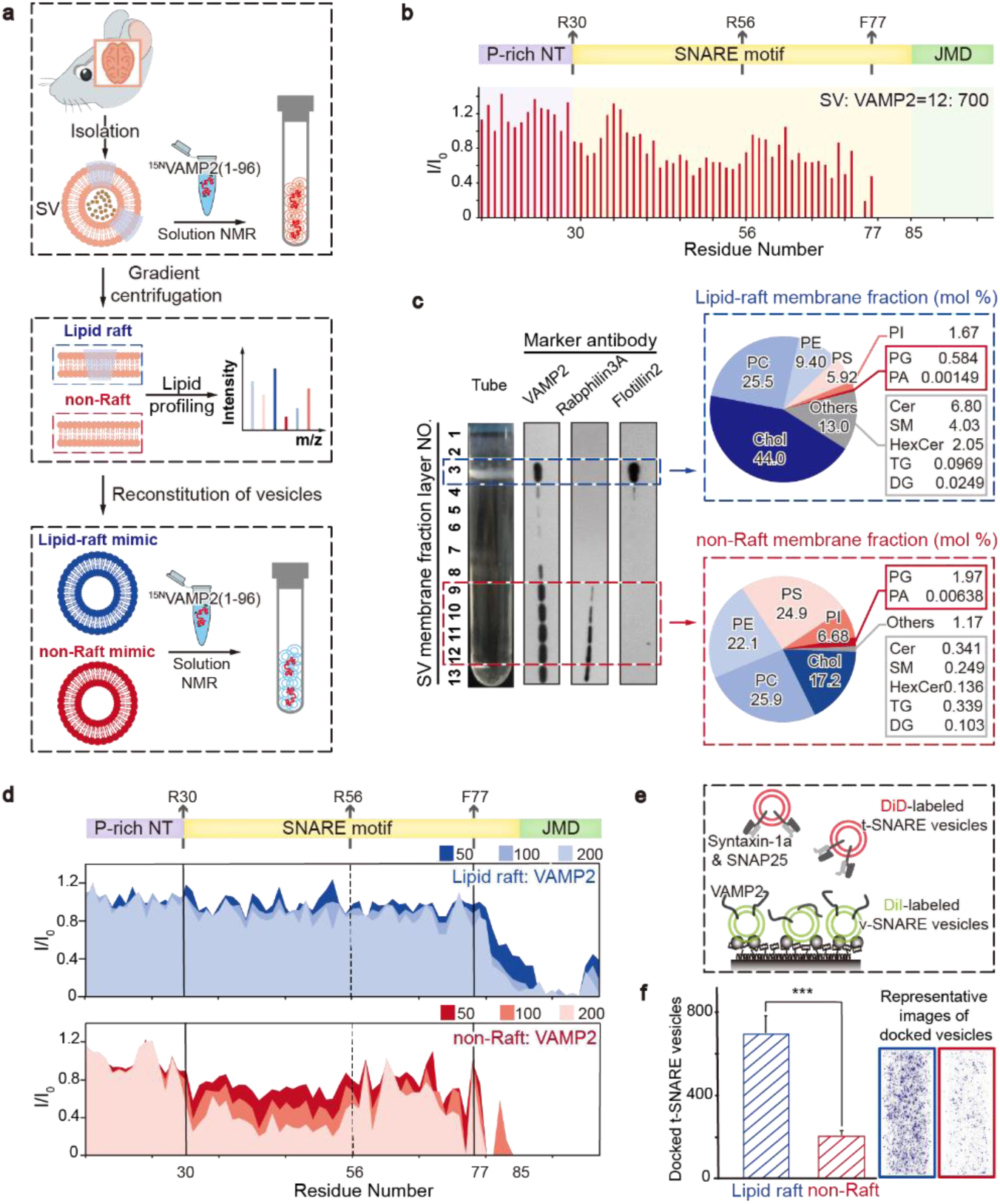
Conformational transition of VAMP2 on membrane sub-domains of synaptic vesicles. **a**, Scheme of experimental design for the structural study of VAMP2 on SV membrane. Upper: SVs isolated from mouse brain were used for NMR titration of VAMP2(1-96) at the physiological ratio. Middle: natural SV membranes were divided into lipid-raft and non-raft membranes by sucrose gradient sedimentation which were analyzed by absolute quantitative lipidomic profiling. Lower: lipid-raft-and non-raft-mimicking vesicles were reconstituted with natural-sourced lipids to titrate VAMP2(1-96). **b**, Residue-resolved NMR signal intensity ratios (I/I_0_) of VAMP2(1-96) titrated by SVs to that in solution. The molar ratio of SV to VAMP2(1-96) is indicated. **c**, Left: distribution of endogenous VAMP2 on SV membrane. SV membranes were fractionated into 13 layers which were collected as lipid raft (layer 3) and non-raft (layers 9-12) according to flotillin2 (lipid raft membrane protein) and rabphilin3A (non-raft membrane protein). Right: lipid compositions of lipid-raft and non-raft membranes by MS-based lipidomic profiling. Chol: cholesterol; PC: phosphatidylcholine; PE: phosphatidylethanolamine; PS: phosphatidylserine; PI: phosphatidylinositol; PG: phosphatidylglycerol; PA: phosphatidic acid; Cer: ceramide; SM: sphingomyelines; HexCer: hexosylceramide; TG: triacylglycerols; DG: diacylglycerols. **d**, Residue-resolved NMR signal intensity ratios (I/I_0_) of VAMP2(1-96) titrated by lipid-raft-mimicking (blue) or non-raft-mimicking (red) vesicles to that in solution at indicated lipid/protein molar ratios. **e**, Scheme of single-vesicle docking assay. A saturated layer of DiI-labeled (green) v-SNARE vesicles carrying full-length VAMP2 was immobilized on the imaging surface. Free DiD-labeled (red) t-SNARE vesicles, reconstituted with syntaxin-1a and SNAP25, were injected into the system. Red laser illumination imaged the t-vesicles that docked on v-vesicles through SNARE complex formation. **f**, Images on the right are representative fluorescence images of the single-vesicle docking assay. The bar graph on the left shows the numbers of lipid-raft-and non-raft-mimicking t-vesicles that docked on v-vesicles. Error bars are standard deviations from 15 random imaging locations in the same sample channel. *** indicates *p*-value < 0.001 by One-way Analysis of Variance (ANOVA) with Turkey Test.

**Fig. 4.**
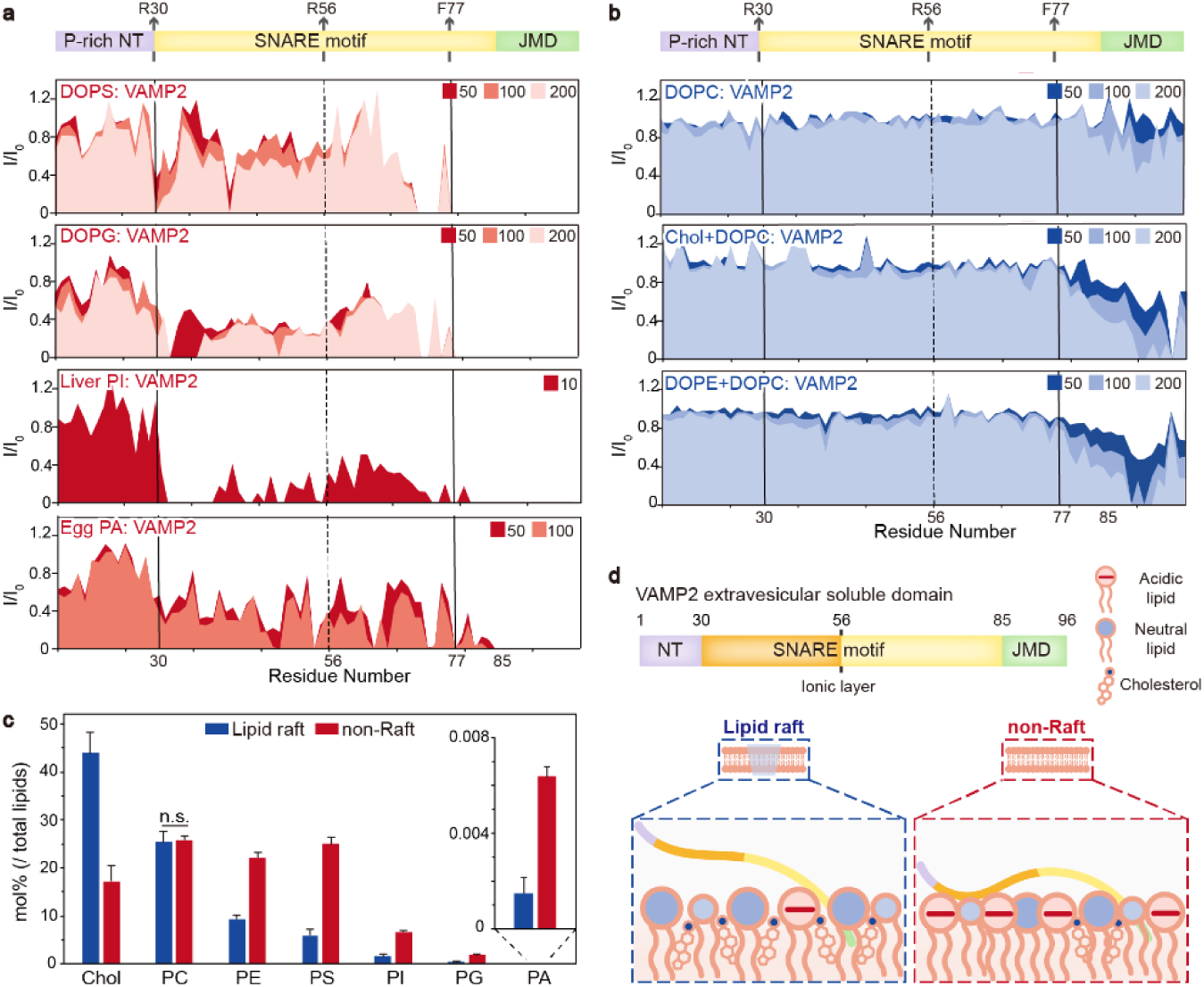
VAMP2 SNARE motif sensors the electrostatic property of membrane surface. **a**, Residue-resolved NMR signal intensity ratios (I/I_0_) of VAMP2(1-96) titrated by acidic liposomes (DOPS, DOPG, liver PI and egg PA) to that in solution at indicated lipid/protein molar ratios. **b**, Residue-resolved NMR signal intensity ratios (I/I_0_) of VAMP2(1-96) titrated by neutral liposomes (DOPC, cholesterol/DOPC (1:1, mol/mol) and DOPE/DOPC (1:1, mol/mol)) to that in solution at indicated lipid/protein molar ratios. **c**, Comparisons of individual lipid classes in lipid-raft and non-raft SV membranes. Error bars are standard deviations from three biological replicates. The data were analyzed by Student’s *t*-test. The *p*-value of all lipid classes, except for PC, are less than 0.001. “n.s.” represents “not significant”. **d**, Schematic presentation of VAMP2 extravesicular domain on lipid-raft and non-raft membranes.

In addition, considering the membrane-binding property of VAMP2, to rule out the possibility that VAMP2 just stuck on the outer surface of cells, rather than entering inside, we conducted the same process of sample preparation yet without electroporation. The 2D ^1^H-^15^N NMR spectrum showed few signals (Supplementary Fig. 5), which confirms that after carefully washing during sample preparation, little VAMP2 remained outside cells, either on cell surface or in solution. We also confirmed that during NMR signal acquirement, no detectable VAMP2 leaked out from cells by immunoblotting (Supplementary Fig. 6). Thus, these results confirm that the in-cell NMR signals are derived from the membrane-associated ^15^N-VAMP2(1-96) in cells.

**Fig. 5.**
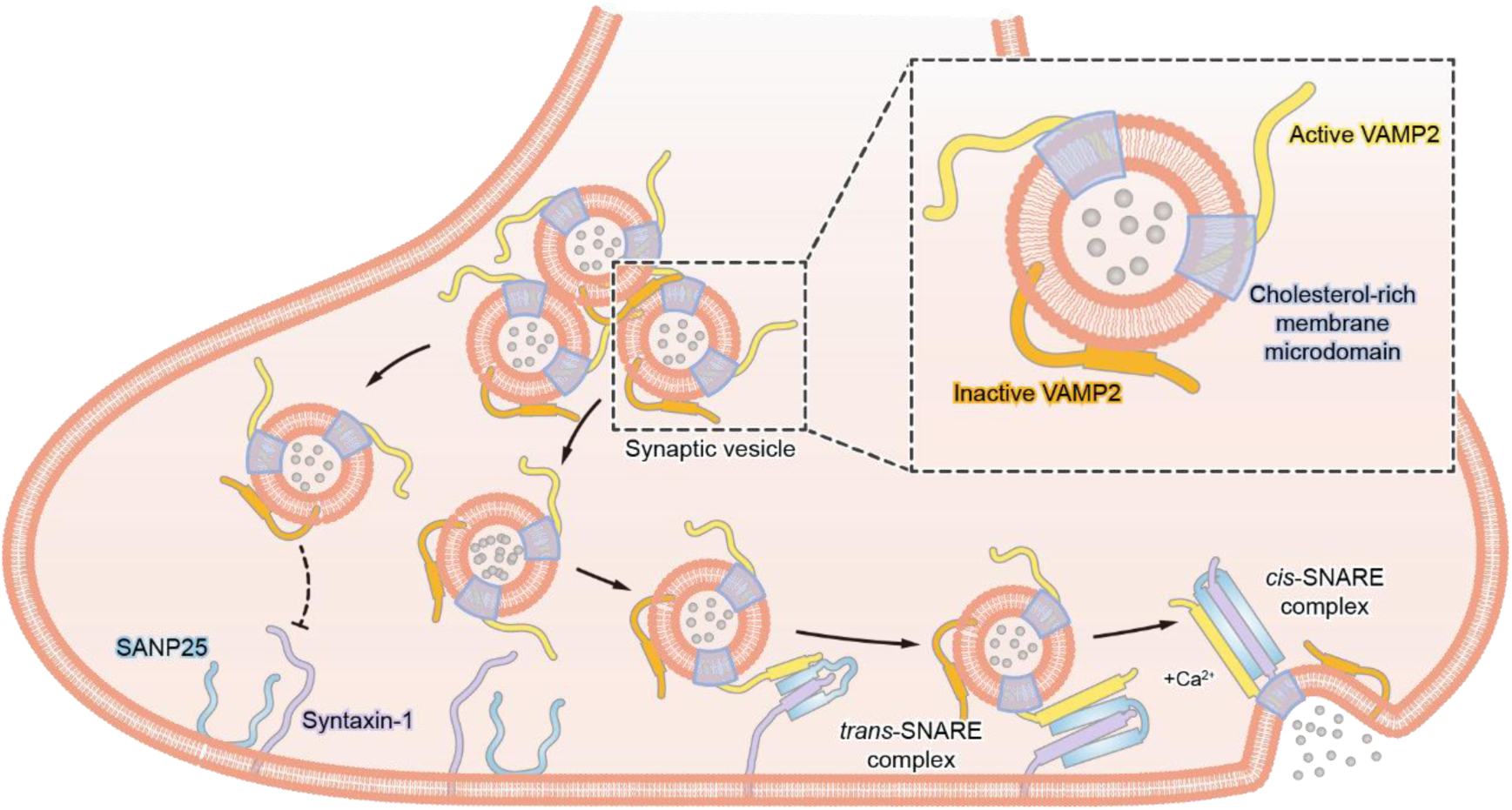
Hypothetic models of VAMP2 configurations in different lipid phases of SV membrane and their engagement in SNARE complex assembly for neurotransmitter release. The extravesicular domain of VAMP2 adopts distinctive conformations in different membrane phases. On the cholesterol-rich microdomains, it is less associated with SV membrane and thus more active to engage in calcium-evoked SNARE assembly and neurotransmitter release. In contrast, in other regions of SV membrane, it tends to hibernate on the membrane with an increased content of α-helical conformation which is relatively inactive in SNARE assembly.

Taken together, these data show that different regions of VAMP2 extravesicular domain exhibit distinctive membrane binding properties: C-terminal residues 78-96 that include the juxta-membrane domain and a short region of SNARE motif, have a strong interaction with cell membranes; the majority of SNARE motif dynamically interacts with membranes; while, the N-terminal proline-rich domain poorly interacts with membranes.

### The conformation of VAMP2 in cells could be regulated by cholesterol

Given the dynamic interaction of VAMP2 SNARE motif with cell membranes, we asked whether the intracellular membrane environment may regulate the membrane association of VAMP2. Previous studies suggested that SANRE proteins are recruited on cholesterol-rich lipid-raft microdomains for SNARE complex assembly^16,17^, thus we manipulated the cholesterol levels of the mammalian cells following a previously established approach^18^to change the intracelluar lipid environment (Fig. 2a). We were able to up-and down-regulate the cellular cholesterol level by ∼ 54% and ∼ 56%, respectively, measured by the absolute quantification using mass spectrometry (MS) (Fig. 2a, Supplementary Fig. 7 and Table 1). We next transported ^15^N-VAMP2(1-96) into cholesterol up-regulated and down-regulated cells by electroporation, respectively (Fig. 2a). Comparable amounts of VAMP2 were delivered into the cholesterol-regulated cells and the untreated cells (Supplementary Fig. 8). Also, cholesterol manipulation did not change the membrane localization of VAMP2 in cells (Supplementary Fig. 8). The 2D ^1^H-^15^N NMR spectra showed that the intensities of VAMP2 signals, but not the chemical shifts, changed in correlation with the intracellular cholesterol levels (Fig. 2b). Notably, as the cholesterol level increased, the intensities of VAMP2 SNARE motif increased as well (Fig. 2c), suggesting a weakened association of the SNARE motif with cell membranes. Conversely, as the cholesterol level decreased, the intensities of SNARE motif decreased (Fig. 2c), suggesting an enhanced membrane association. In contrast, the flexible N-terminal domain of VAMP2 showed arbitrary intensity changes in cholesterol manipulated cells (Fig. 2c).

### VAMP2 exhibits distinctive conformations and activities in different lipid phases of SV membrane

Given that cholesterols could alter the lipid distribution of cell membranes^19^, the results derived from in-cell NMR experiments evoked us to speculate that the conformation of VAMP2 may vary in different lipid phases of SV membranes. Thus, we first isolated SVs from mouse brains, and then titrated ^15^N-VAMP2(1-96) with mouse SVs (Fig. 3a, upper). The molar ratio of VAMP2 to SV was 700: 12 which is close to that within the synaptic bouton^20^. The result showed that the 2D NMR spectrum of VAMP2 with SVs exhibited a general similarity to that of VAMP2 in cells (Supplementary Fig. 9 and Fig. 3b). Then, we fractionated SV membranes into cholesterol-enriched lipid-raft and non-raft fractions by sucrose gradient centrifugation. (Fig. 3a, middle, and Fig. 3c, left). Immunoblot showed that the endogenous VAMP2 presented in both the lipid-raft and non-raft fractions (Fig. 3c, left).

Next, to investigate the configurations of VAMP2 in these two different membrane phases, we first quantitatively characterized the lipid compositions of the lipid-raft and non-raft fractions by using MS-based lipidomic profiling (Fig. 2a, middle, Fig. 2c and Supplementary Table 2). The result showed that both lipid-raft and non-raft membranes are dominantly composed of glycerol-phospholipids (including PC, PE, PS, PI, PG and PA) and cholesterol. Based on the lipidomic analyses of SV membrane subdomains, we used natural-source cholesterol and glycerol-phospholipids to reconstruct vesicles mimicking the lipid compositions of the lipid-raft and non-raft membranes, respectively (Supplementary Fig. 10). Then, we titrated them to ^15^N-VAMP2(1-96) and found a remarkable difference as VAMP2 binds to these two different membranes (Fig. 3a, lower, Fig. 3d and Supplementary Fig. 11). Overall, VAMP2 bound much tighter with the non-raft vesicle than with the lipid-raft vesicle. As VAMP2 binds to the non-raft vesicle, the SNARE motif, especially the N-terminal half of the motif, exhibited an enhanced interaction with membrane as the concentration of non-raft vesicle increased (Fig. 3d, lower). In contrast, as VAMP2 binds to the lipid-raft vesicle, the SNARE motif of VAMP2 barely interacted with membranes (Fig. 3d, upper).

We next asked whether the conformational difference of VAMP2 on membrane lipid raft and non-raft influences the SNARE assembly. Thus, we reconstituted full-length VAMP2 on lipid-raft or non-raft vesicles as v-SNARE vesicles, and reconstituted syntaxin-1a and SNAP25 on lipid-raft or non-raft vesicles as t-SNARE vesicles (Fig. 3e). Stable assembly of these three proteins into the SNARE complex leads to the docking of a t-vesicle to the immobilized v-vesicle, which could be monitored by the single-vesicle fluorescence microscopy. The result showed that the v-vesicle mimicking lipid raft presented a significantly higher docking efficiency than that mimicking the non-raft (Fig. 3f).

Altogether, these results demonstrate that VAMP2, especially the SNARE motif, exhibits distinctive configurations in different lipid phases of SV membrane. As localizing on the cholesterol-rich lipid rafts, the SNARE motif is free of membrane binding and readily to associate with its homologous motifs of other core SNARE proteins. While, as localizing in the non-raft phase of SV membrane, the SNARE motif is more associated with membrane and less active for SNARE assembly.

### The membrane binding of VAMP2 is strengthened by acidic lipids

To investigate the mechanism underlying the different conformations of VAMP2 in different membrane phases, we tested the individual lipid components of SV membrane for their influence on the structure of VAMP2. The result of lipid analysis showed that SV membranes mainly consist of cholesterol and glycol-phospholipids including neutral lipids (PC and PE) and acidic lipids (PS, PI, PG and PA) (Fig. 3c). Thus, we titrated VAMP2(1-96) with each individual lipid and monitored their interactions with VAMP2. The analyzed NMR data showed that VAMP2 exhibited strong interactions with the negatively-charged phospholipids (Fig. 4a). A regional binding was also observed for the binding of VAMP2 to negatively-charged lipids, which is similar to that observed in cells and in the binding of VAMP2 to SVs (Fig. 4a, 1g, and 3b). In contrast, except for the juxta-membrane domain, VAMP2 showed no significant interaction with cholesterol or the neutral phospholipids (Fig. 4b). In addition, lipid profiling showed that the cholesterol-rich lipid rafts contain significantly less negatively-charged lipids than the non-raft membranes (Fig. 4c). Thus, these results indicate that the conformation of VAMP2 on different lipid phases of SV membrane could be defined by the electrostatic properties of the membrane surface due to different lipid compositions (Fig. 4d).

## DISCUSSION

In-cell NMR spectroscopy is a cutting-edge technology that provides residue-specific structural information of proteins in live cells^21^. In this work, we were able to perform in-cell NMR to study a membrane-associated protein — VAMP2, and captured its structural changes upon the lipid environment changes in mammalian cells. Combined with solution NMR and MS-based lipidomic profiling, we further demonstrated that the SNARE motif configures differently in different lipid phases of SV membrane (Fig. 5). When localizing on the cholesterol-rich lipid rafts, the SNARE motif of VAMP2 is less associated with membranes and more active to engage in SNARE assembly with syntaxin-1 and SNAP25 (Fig. 5). When localizing elsewhere on SV membranes, the SNARE motif is more associated with membranes probably with an α-helical conformation^10,11^, and is less active for SNARE assembly (Fig. 5).

Membrane microdomains have been suggested as functional loci of various proteins involved in different pathways^22^. Previous studies show that the SANRE proteins are highly enriched on lipid rafts in different cells^16,17^. Reducing the cellular cholesterol level to disrupt lipid rafts can impair the regulated exocytosis^17^. In addition, in the Niemann-Pick disease type C1, cholesterol is deficient to be recruited to neuronal membranes, and thus shows dramatic defects in evoked neurotransmission^23^. Our work indicates that membrane microdomains not only provide a place to recruit function-linked proteins, and also may directly configure the proteins to facilitate protein interactions. Therefore, such defects of exocytosis may because of more non-lipid raft phases of membrane in cholesterol deficient cells where VAMPs hardly assemble with other SNAREs. Both membrane lipids and proteins (e.g. syntaxin-1^24^and synaptotagmin-1^25^) may cooperate to regulate VAMP2 structure, but lipid distribution and phase separation may add a new spatial dimension to generally modulate SNARE protein conformations and SNARE complex assembly.

In this work, we quantitatively profiled the lipid compositions of lipid-raft and non-raft phases of SV membrane (Supplementary Table 2). We have shown that the different electrostatic surfaces of these two phases lead to their different binding affinities to the SNARE motif of VAMP2. Actually, we also noticed differences between lipid rafts and non-rafts in the length and saturation of lipid aliphatic tails. Specifically, phospholipids with short tails are enriched in the lipid rafts (Supplementary Fig. 12a) which are easy to flip between inner and outer membrane leaflets^26^. This is consistent with the knowledge that evoked vesicle fusion requires fast lipid flipping^27^. While, phospholipids with unsaturated tails are enriched in the non-raft membranes (Supplementary Fig. 12b). A previous study reported that high contents of unsaturated fatty acyl chains can help to support rapid vesicle fission because of its low energetic cost in membrane bending^28^. Combining this report with our MS data may probably explain the phenomenon that cholesterol-depleted neurons, non-lipid raft increase, have an increased level of spontaneous fusion^29^.

The extravesicular domain of VAMP2 features an intrinsically disordered sequence which is unstructured in the form of monomer. Proteins containing intrinsically disordered regions (IDRs) present as a heterogeneous conformational ensemble, which may undergo distinct conformational changes in different biological contexts. As for VAMP2, we demonstrate that it transforms between different conformations as localizing on different regions of SV membranes and in complex with other core SNARE proteins. About 70% of human membrane proteins involved in signaling contain IDRs^30^. Many of them (e.g. T cell receptor-CD3^31^and epidermal growth factor EGFR^32^) have multiple binding partners including proteins, lipids and metabolic ions. The technologies performed in this study may be useful for the structural study of other IDR-containing membrane proteins in live cells and physiological-relevant circumstances.

## Acknowledgments

We thank Songzi Jiang and other staff members of National Facility for Protein Science in Shanghai for assistance in NMR data collection. We thank Jinshi Shen (Colorado, U.S.A.) and Chenqi Xu (Shanghai Institutes for Biological Sciences, CAS.) for presenting proteins and the serum to us.

## Data and materials availability

All data is available in the main text or the supplementary materials.

## Supplementary Information

Materials and Methods

References (33-44)

Supplementary Figures 1-12

Supplementary Table 1-2 (separate Excel file), 3

