## Supplementary information for "Membrane association of VAMP2 SNARE motif in cells and its regulation by different lipid phases of synaptic vesicle membrane"

**This file includes:**

Materials and Methods: *page 2-7*

References: *page 8*

Supplementary Figure. 1 to 12: *page 9-20*

Supplementary Table 1-2 (separate Excel file) & 3: *page 21*

Materials and Methods

**Lipids**

The lipid molecules used in this study were purchased from Avanti Polar Lipids as follows: Cholesterol(ovine) (700000), DOPC (850357), DOPE (850725), DOPS (840035), DOPG (840475), Brain PC (840053), Brain PE (840022), Brain PS (840032), Egg PG (841138), Liver PI (840042), Egg PA (840101), Biotinyl PE (870828).

**Protein purification**

The rat VAMP2(1-96), VAMP2(1-78) and VAMP2(1-59) fused with His-tags were overexpressed from pET31b plasmids in *E. coli* BL21-DE3 (CodenPlus). Non-isotope enriched proteins were produced in LB medium, ^15^N-labeled proteins were produced in M9 minimal media supplemented with ^15^NH_4_Cl (1 g/L, CIL). Bacteria were harvested by centrifugation after induction by 1 mM IPTG at 37 °C for 6 h in LB medium or 12 h in M9 media with the OD_600_ value of around 2.0. The bacteria were lysed by high pressure in a lysis buffer (50 mM Tris-HCl, pH 7.4, 100 mM NaCl, 1 mM PMSF).

Since the overexpressed VAMP2(1-96) forms inclusion bodies *in E.coli* the inclusion bodies in pellets were spun down (16,000 g for 30 min) and washed twice in 10% Trinton-X100 and 1M NaCl buffer (25 mM Tris-HCl, pH 7.4) to remove lipids, nucleotides and other proteins. Then the inclusion bodies were solubilized in 6M guanidine hydrochloride buffer (25 mM Tris-HCl, pH 7.4) and purified by HisTrap^TM^ HP columns (GE Healthcare). The purified protein was harvested in 6M guanidine hydrochloride buffer (25 mM sodium phosphate, pH 2.0) and further purified by RP-HPLC C3 column (Agilent Technology). The lyophilized protein was solubilized and cleaved by TEV protease at 4 °C overnight in buffer (50 mM Tris-HCl, pH 7.4, 100 mM NaCl, 1mM DTT) to remove His-tag. Finally, the VAMP2(1-96) proteins without His-tag was purified by RP-HPLC C8 column and lyophilized.

VAMP2(1-78) and VAMP2(1-59) were overexpressed in *E.coli* as soluble proteins in the supernatants (after centrifugation at 16,000 g for 30 min) and were purified by HisTrap^TM^ HP columns. Then, the His-tag proteins were dialyzed into the cleavage buffer as above and cleaved by TEV protease at 4 °C overnight. At last, the enzyme-digested VAMP2 proteins were purified by RP-HPLC C8 column and lyophilized.

The rat full-length VAMP2, syntaxin-1a and SNAP25 were gifted from lab of Jinshi Shen (Colorado, U.S.A.). The three proteins were purified as previously reported (*33*). The human α-synuclein protein was purified by following the protocol previously published (*34*).

**Electroporation of purified proteins into mammalian cells**

Human HEK-293T (ATCC, CRL-3216) and SH-SY5Y (ATCC, CRL-2266) cells were cultured following the protocol provided by ATCC. Both cell lines were tested for mycoplasma contaminations and were mycoplasma free. The cells with four to six passage were used for NMR experiments.

The purified protein powder of VAMP2(1-96), VAMP2(1-78), VAMP2(1-59) or α-synuclein was dissolved in Buffer R supplied in the Neon transfection system kit (Invitrogen, MPK10025) to a final concentration of 300 μM. Cells were collected by trypsinization and washed with PBS for three times to remove the culture medium. Then the cells were resuspended with VAMP2 solution with the density of 8×10^7^ cells/mL for HEK-293T and 4×10^7^ cells/mL for SH-SY5Y. Electroporation was conducted using 100 μL of the cells mixed with protein with a pulse program of 1,400 V (pulse voltage), 20 ms (pulse width) and 2 pulses by the Neon transfection system (Invitrogen, MPK5000). The control sample was conducted with the identical setup but without the electric shocks.

For immunofluorescence and time-course immunoblotting experiments, aliquots of 0.5×10^6^ cells were added to each well in a 24-well plate, filled with 0.5 mL medium. For in-cell NMR and sub-cellular fractionation experiments, aliquots of 4~8×10^6^ cells were added to eight 10-cm dishes with 10 mL medium and cultured for 3~6 h for cell recovery. Then the cells were harvested and washed with PBS for four times. The in-cell NMR sample in 160 μL pH-stable L-15 medium (Gibco, 11415064) and 40 μL D_2_O was settled into the NMR tube by gentile sedimentation with a hand-cranked centrifuge. The suspended cells and additional medium were discarded. Finally, 500 μL sedimented cell slurry was prepared for NMR measurement. The cell sample for sub-cellular fractionation were stored at -80°C prior to the experiments.

To determine the intracellular concentration of VAMP2(1-96), and check its potential cell leakage after in-cell NMR experiments, the cell samples were centrifugated at 300 g for 3 min. Then the the supernatants (S) and pellets (P) were resuspended in Laemmli buffer to volume of 500 μL, and boiled for 10 min. As for the immunoblotting experiment, the samples were diluted by 10-times and 10 μL sample was loaded in each lane of a 15% SDS-PAGE gel. The concentration of the electroporated VAMP2(1-96) in-HEK-293T-cell (~10 μM) and in-SH-SY5Y-cell (~80 μM) samples were analyzed and calculated by using ImageJ (*35*) (Fig. 1c).

**Sub-cellular fractionation of the electroporated cells**

Isolation of the cytosol and membrane fractions of the cells was achieved by following a previously published protocol (*36*). The experiment was performed at 4°C or on ice with pre-cooled reagents. Briefly, aliquots of 2~4×10^7^ electroporated cells were permeabilized by adding 2 mL of digitonin lysis buffer (110 mM KOAc, 2mM MgCl_2_, 20 mM K-HEPES, pH 7.5, 0.02% digitonin) with protease inhibitors for 10 min to release cytosolic contents. The lysate (Lys) was centrifuged at 20,000 g for 10 min. The supernatant was the cytosol fraction (Cyto). The permeabilized cell pellet was washed three times by the lysis buffer and centrifuged at 20,000 g for 10 min. The pellet was resuspended in 2 mL lysis buffer as for the membrane fractions (Mem).

The Lys, Cyto and Mem samples were further characterized by immunoblotting. The antibodies used include VAMP2 (SYSY, 104211, 1: 10,000), GAPDH (Cell Signaling Technology, 2118, 1:1,000) for cytosol, IRE1α (Cell Signaling Technology, 14C10, 1:1,000) for endoplasmic reticulum, ACSL4 (Santa Cruz, sc-365230, 1: 1,000) for plasma membrane-associated membranes, and VDAC (Cell Signaling Technology, 4661, 1:1,000) for mitochondria. Immunoblotting quantification (Fig. 1d, and Supplementary Fig. 4a) of the proportion of electroporated protein in cytosol (Cyto/(Cyto+Mem)) was analyzed by ImageJ (*35*).

**Immunofluorescence staining of the cultured cells**

For immunofluorescence imaging, cells with or without the electroporated proteins were recovered for 3~6 h on poly-D-lysine-coated coverslips in 24-well plates. Then, the cells were washed by pre-warmed PBS three times to remove extracellular un-delivered proteins, then fixed in 4% (w/v) paraformaldehyde in PBS for 30 min and permeabilized with 0.1% (v/v) Triton X-100 in PBS for 20 min. After washing with PBS three times, cells were blocked with 10% BSA in PBS for 1 h followed by incubation of antibodies at 4 °C overnight, including Oyster-550 labeled VAMP2 (SYSY, 104211C3, 1: 1,000), α-synuclein (BD Biosciences, 610787, 1: 1,000) and FITC-labeled phalloidin (Yeasen, 40736ES75, 1:200) for F-actin filaments beneath cell membranes (*37*). Then Alexa Fluor 594-conjugated secondary antibodies (Invitrogen, A-11020, 1:500) were used. Slides were then washed with PBS for three times. The nucleus was stained by antifade mountant coupled DAPI (Invitrogen, P36935). Finally, the samples were observed by a confocal microscope (Lecia, SP8).

**In-cell and *in vitro* solution NMR spectroscopy**

All the NMR experiments were carried out at 25 °C on a Bruker 900 MHz spectrometer equipped with a cryogenic probe. The buffer used in all of the *in vitro* NMR experiment was 50 mM sodium phosphate buffer (pH 6.5) containing 50 mM NaCl and 10% D_2_O (v/v). For the *in vitro* titration experiments, liposomes were prepared with a concentration of 50 mM, and were gradually added into the solution containing 25 μM ^15^N-VAMP2(1-96) to the indicated molar ratios with a final volume of 500 μL for NMR measurement. As for the SV titration experiment, the concentration of SV was calculated according to the ratio of its total proteins and phospholipid which was previously used (*20*). Bruker standard SOFAST-HMQC pulse sequence (*38, 39*) was used and the ^1^H shape pulse efficient was optimized for collecting the 2D NMR spectrum of the in-cell samples with 80 scans. The delay time (D1) was set to 0.29 s, and 1024 and 128 complex points were used for ^1^H and ^15^N, respectively.

The backbone ^15^N relaxation parameters of R_1_, R_2_ were recorded from the VAMP2 electroporated HEK-292T cell samples as well as 50 μM VAMP2(1-96) in-solution protein sample, respectively. For in-solution NMR samples, the time delays for R_1_ experiment were 10, 40, 100, 200, 300, 500, 700, 800, 1200, and 2000 ms, while those for R_2_ experiments were 0, 20, 40, 80, 120, 200, and 400 ms. For in-cell NMR samples, the time delays for R_1_ experiment were 10, 40, 100, 200, 300, 500, 700, 900, and 1200 ms, while those for R_2_ experiments were 0, 20, 50, 80, 120, 200 and 400ms.

Backbone resonance assignment of VAMP2(1-96) was accomplished according to the previously published assignments (BMRB 4272) (*8*). Residue T27, S61 and eight prolines in the N-terminal of VAMP2 cannot be assigned. All of the NMR data were processed by NMRpipe (*40*) and analyzed by SPARKY (*41*). Specifically, the R_1_ and R_2_ relaxation data were analyzed using SPAKY relaxation fitting extension. The Residue-resolved relative signal intensity ratios (Y) of in-cell (I) to in-solution (I_0_) NMR spectrum were calculated for each residue X as following equation: Y=[I(X)/I_0_(X)]/[I(5)/I_0_(5)], where the intensity ratio of the residue X was normalized by the highly flexible residue 5 for comparing the regional flexibility of VAMP2(1-96) in cells (Fig. 1f). The residue-resolved relative NMR signal changes (Z) of VAMP2(1-96) in different treated cells (Y) comparing to it in untreated control cells (Y_0_) were calculated as following equation: Z(X)= (Y-Y_0_)/Y_0_ (Fig. 2c).

**Manipulation of the cellular cholesterol level**

Manipulation of the cholesterol level within cells was achieved by previous protocols (*18*). In brief, to elevate the cellular cholesterol level, cells were treated with 20 μg/mL Cholesterol-MβCD (Sigma, C4951) for 8 h before the electroporation experiment and lipidomic profiling. To decrease the cellular cholesterol level, the cell cultured medium was replaced to lipoprotein deficient serum (LPDS) supplemented medium containing 5 μM mevastatin and 50 μM mevalonate (Sigma, M2537 & 90469) and the cells were further cultured for 8 h. Then the cells were treated with 5 mM MβCD for 20 min prior to the electroporation experiment and lipidomic profiling. The LPDS was gifted from lab of Chenqi Xu (Shanghai Institutes for Biological Sciences, CAS.), which was prepared as previously published protocol (*18*).

**Isolation of synaptic vesicles from mouse brains**

Synaptic vesicles (SV) were purified following a previously published protocol (*42*). The experiment was performed at 4°C or on ice with pre-cooled reagents. Four brains from 8-week-old C57BL6 mice (male) were homogenized (900 r.p.m. for 10 min) in 25 mL 4 mM Na-HEPES, pH 7.4 and 320 mM sucrose buffer (HB) with protease inhibitors by using a PTFE pestle in a 40 mL glass tube (Sigma, P7984). The homogenate was centrifuged at 1,500 g for 10 min. The supernatant (S1) was collected and kept on ice. The pellet was resuspended with 25 mL HB and homogenized (900 r.p.m. for 10 min), followed by centrifugation at 1,500 g for 10 min. Then the supernatant (S2) combined with S1 were centrifuged at 20,000 g for 20 min. The pellet which contains synaptosomes was resuspended with 2 mL HB and then homogenized in 20 mL H_2_O at 1,200 r.p.m. for 10 min followed by adding 50 μL of 1M Na-HEPES, pH 7.4 and protease inhibitors. The homogenate was placed on ice for 30 min, and centrifuged at 20,000 g for 20min. The supernatant was ultra-centrifuged at 70,000 g for 45 min. The pellet was clustered SV extracts. The samples for different fraction were diluted with a total protein concentration of 50 μg/mL for immunoblotting. Antibodies used included synaptophysin (Sigma, S5768, 1:1,000) for SVs, VDAC for mitochondria, IRE1α for endoplasmic reticulum and GAPDH for cytoplasm. To obtain homogenous SVs, fraction SV was resuspended in NMR buffer and homogenized by using a PTFE pestle in a 3 mL glass tube (Sigma, P7734) at 1,200 r.p.m. for 10 min. Furthermore, to disrupt any remaining SV clusters before NMR experiment, the homogenized SVs was drawn through a 20-gauge hypodermic needle attached to a 10-ml syringe, and then changed to a 27-gauge needle and expelled.

**Isolation of lipid-raft and non-raft membranes from SVs**

Fractionation of lipid-raft and non-raft membranes from SVs was achieved by using a previously published protocol (*43*). Briefly, the isolated SV was suspended with 4 mL ice-cold Mes-buffer (25 mM Mes, pH 6.5, 150 mM NaCl, phosphatase inhibitors) supplemented with Triton X-100 to a final concentration of 1% v/v. The mixture was gently swung at 4 °C for 10 min, and then homogenized using a PTFE pestle in an 8 mL glass tube (Sigma, P7859) at 200 r.p.m. for 10 min, followed by adding 80% (w/v) sucrose in Mes-buffer with a final concentration of sucrose of 40% (w/v). The mixture was divided equally into two 13.5 mL ultracentrifuge tubes (Beckman, 344059) and overlaid successively with 6 mL 30% (w/v) sucrose and 2.5 mL 5% (w/v) sucrose. After centrifugation (Beckman, SW41Ti rotor) at 240,000 g for 6 h, 12 fractions (1 mL/fraction) were collected from the top to the bottom. The pellet was resuspended in Mes-buffer to a total volume of 1 mL as fraction 13. The different fractions were analyzed by immunoblotting. Antibodies used in the experiment included flotillin2 (Santa Cruz, sc-28320, 1:500) for lipid rafts, rabphilin3A (Santa Cruz, sc-393197, 1:500) for detergent-soluble non-raft membranes and VAMP2. By analysis of the immunoblot, the fraction 3 was stored as the lipid-raft sample and fraction 9-12 were mixed stored as the non-raft sample.

**Lipid extraction from biological samples**

The lipids in cells, lipid raft and non-raft fractions were extracted using a modified MTBE extraction method. 200 μL of each sample was mixed with 480 μL extraction solvent (MTBE: MeOH = 5:1, v/v). The sample was vortexed for 30 s, followed by 10 min sonication and 15 min centrifugation at 3,000 r.p.m. The upper organic layer (i.e., MTBE layer) was collected. Then, 200 μL of MTBE was added to the left aqueous layer for re-extraction. The re-extraction process was repeated twice, and the pooled organic layer was evaporated using a vacuum concentrator. The dried extract was resolved in 100 μL of DCM: MeOH (1:1, v/v) containing 2 μL of a stable isotope labeled internal standard (IS, d7-PE (15:0/18:1), 100 ppm). The total protein concentration of SV and cell samples for lipidomic profiling were measured by a Pierce BCA Protein Assay Kit (Thermo Fisher Scientific, 23225).

**LC-MS/MS analysis for lipidomic profiling**

The LC-MS/MS analysis were performed by using a UHPLC system (Agilent Technologies, 1290 series) coupled to a quadrupole time-of-flight mass spectrometer (Sciex, TripleTOF 6600). Chromatographic separations were performed on Phenomenex Kinetex C18 column (particle size, 1.7 μm; 100 mm (length)×2.1 mm (i.d.)) with a column temperature of 55 °C. The mobile phases A = 10 mM ammonium formate in H_2_O: ACN (6:4, v/v), and B = 10 mM ammonium formate in IPA: ACN (9:1, v/v), were used for both ESI positive and negative modes. The linear gradient elutes from 40 to 100% B (0–12min), 100% B (12–14min), 100 to 40% B (14–14.2 min), then equilibrate at 40 % B until 18 min. And the flow rate was set as 0.3 mL/min. The mass spectrometry parameters were applied as follows: ion source gas 1 (GS1), 60 psi; ion source gas 2 (GS2), 60 psi; curtain gas (CUR), 30 psi; temperature, 600 °C; ion-spray voltage floating (ISVF), 5000 V or -4500 V in positive or negative modes, respectively; de-clustering potential (DP), 100 V. Data-dependent acquisition (DDA) method was used for MS/MS acquisition. Each acquisition cycle consists of one rapid TOF MS survey scan (200 ms) followed by the consecutive acquisition of 11 product ions scans (50 ms each). For untargeted lipidomic analysis, a data-dependent acquisition (DDA) method is used for data acquisition, in which, one cycle consists of one rapid TOF MS survey scan (200 ms) followed by the consecutive acquisition of 11 product ion scan (50 ms per spectrum). The collision energy (CE) was set as 45 V, and CE spread was set as 15V, for both positive and negative modes.

**Lipid identification and absolute quantification**

LC-MS raw data (.wiff) files were converted to the mzXML format using ProteoWizard 3.0.6150, and processed by LipidAnlayzer (*44*). Briefly, peak detection and alignment were performed by the CentWave algorithm and the ordered bijective interpolated warping (OBI-Warp) algorithm. The acquired MS/MS spectra were identified through the spectral match using an in-house MS/MS spectral library. A similarity score was calculated using the dot product function. The similarity score ranges from 0 to 1, referring to no similarity and a perfect match, respectively. Lipid matches with scores larger than 0.8 were kept as candidates, and the top hits were recognized as identification. Finally, the absolute quantification of lipids can be achieved through response factor-based approach using the peak area and internal standard information. For absolute quantification of cholesterol, an external standard curve of cholesterol was measured.

**Preparation of liposomes and fluorescent labeled vesicles reconstituted with SNAREs**

The different lipid molecules for liposome preparation used in this study were dissolved and mixed in chloroform and evaporated using a dry nitrogen stream. The dried lipid film was hydrated by adding the NMR buffer or HEPES buffer (100 mM NaCl, 25 mM HEPES, pH 7.4), and then ultrasonicated in a water bath at 65 °C for 10 min. To prepare liposomes with homogenous size, the hydrated lipids were extruded 21 times at 65 °C through a polycarbonate film with a pore size of 50 nm (Whatman Nucleopore Track-Etch) by using an extruder apparatus (Avanti Polar Lipids, 610000). The size and homogeneity of the liposomes were confirmed by dynamic light scattering instrument (Wyatt Technology, 431-DPN).

In single-vesicle clustering experiment, to prepare the SNARE protein reconstituted labeled vesicles, 2 mol% of brain PC and 0.5 mol% of brain PE of the lipid-raft and non-raft vesicles were substituted by 2 mol% DiI or DiD (Invitrogen, D282 or D307) and 0.5 mol% biotinyl PE. Then, DiI labeled vesicles and full-length VAMP2 were mixed and incubated for 30 min on ice. DiD labeled vesicles were added to pre-mixed syntaxin-1a and SNAP25 solution. Both of the mixtures were diluted with the same volume of HEPES buffer and dialysis in 2 L HEPES buffer overnight, the reconstituted vesicles were transferred into tubes for further research. The lipid-to-protein molar ratio was 200:1 based on VAMP2 or syntaxin-1a.

**Single-vesicle clustering experiments**

The prepared DiI and DiD vesicles reconstituted with neuronal SNARE proteins were used for single-vesicle clustering experiments. The lipid to protein ratio was 200:1. In brief, the DiI vesicles reconstituted with VAMP2 were firstly immobilized on the imaging surface of PEGylated quartz slides via a biotin/NeutrAvidin interaction. The DiI vesicle coverage was confirmed by a green laser excitation. After buffer change, DiD vesicles harboring syntaxin-1a & SNAP25 were injected into the sample channel and incubated for 30 min to be bound to the surface of immobilized v-SNARE vesicles. Before imaging under a wide-field total internal reflection fluorescence (TIRF) microscopy, the channels were washed by HEPES buffer three times to remove uncombined vesicles.

The smCamera program was used to acquire and analyze the images respectively. The number of vesicles clustering was determined via counting the number of fluorescent spots of acceptor channel under red laser excitation. 15 random locations were imaged and analyzed for each sample channel on the slide. The compared results were expressed as mean ± standard deviations. One-way Analysis of Variance (ANOVA) with Turkey Test was used to determine the statistical significance among different groups. When P﹤0.001, the statistics were considered extremely significant (***).

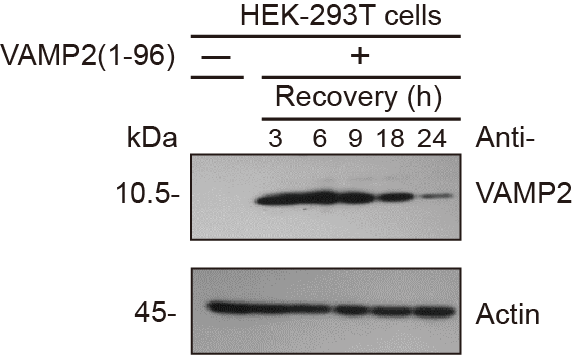
**Supplementary Fig. 1 | Half-life of delivered VAMP2 in cells.** The relative amounts of VAMP2(1-96) in cells were measured by immunoblotting at different recovering time.

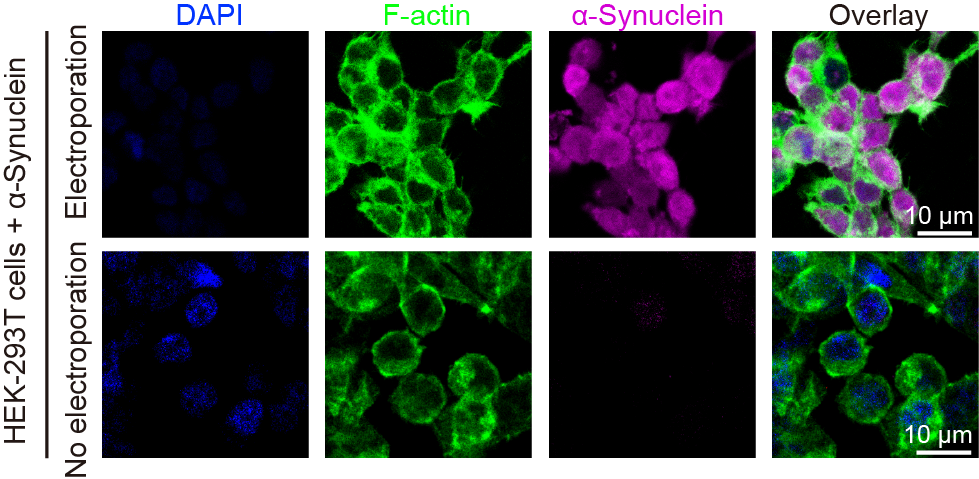
 Supplementary Fig. 2 | Distribution of delivered α-synuclein in cells. Cellular localization of α-synuclein was visualized by immunofluorescence staining. F-actin filaments depict cell skeletons stained by FITC-phalloidin and nuclei were stained by DAPI.

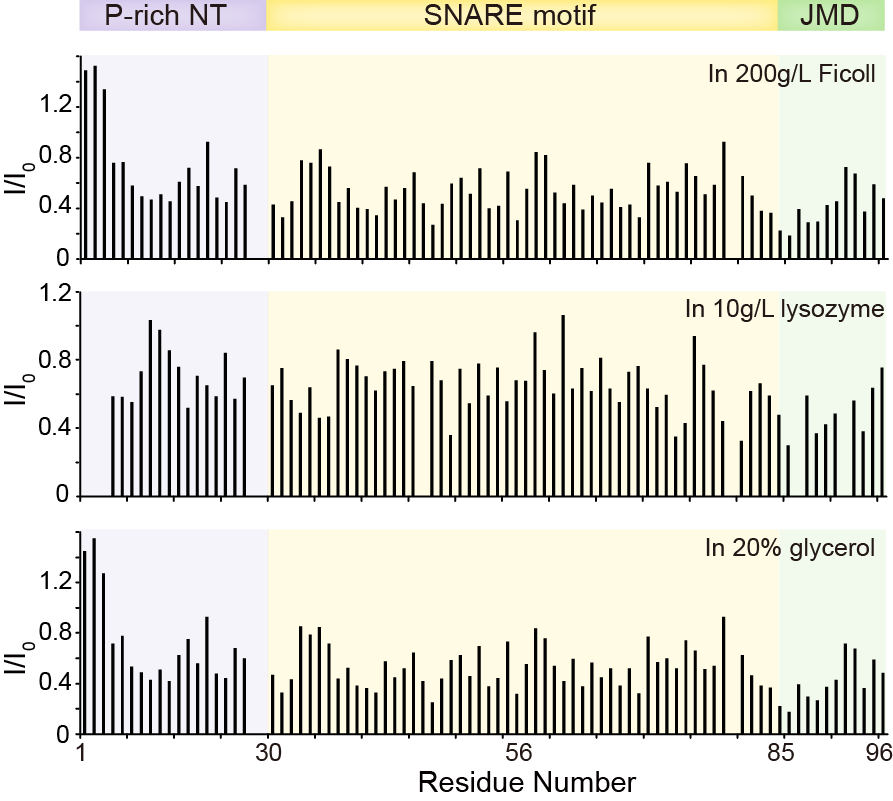
 Supplementary Fig. 3 | NMR titration of VAMP2 with different crowding agents. The concentration of VAMP2(1-96) is 25 μM. Concentrations of the crowding agents are indicated.

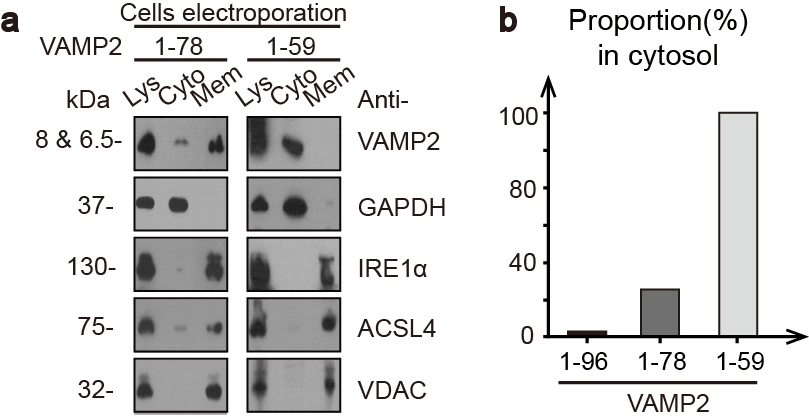
 Supplementary Fig. 4 | Sub-cellular localization of VAMP2 variants. a, Localization of delivered VAMP2(1-78) and VAMP2(1-59) in HEK-293T cells by immunoblotting. Fractions of the total lysates (Lys), cytosol (Cyto) and membrane (Mem) were validated by immunoblotting with antibodies noted in Fig. 1D. b, Quantification of different VAMP2 variants in cytosol based on (A) and Fig. 1D. Detailed information of data processing was noted in the Supplementary Methods.

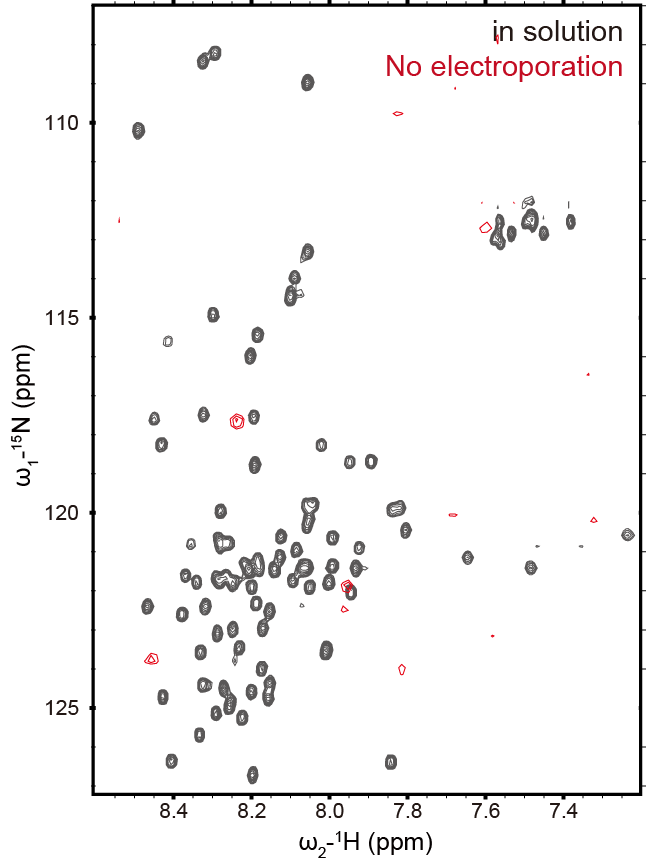
 Supplementary Fig. 5 | VAMP2 remainder on cell outer surface detected by NMR spectroscopy. HEK-293T cells were treated with VAMP2(1-96) in the same way as the sample preparation for in-cell NMR except for electroporation.

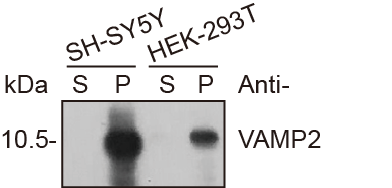
 Supplementary Fig. 6 | Leakage of VAMP2 from cells after NMR measurement. The supernatant medium (S) and cell pellet (P) were collected for checking the leakage of VAMP2(1-96) from cells over NMR measurement by immunoblotting.

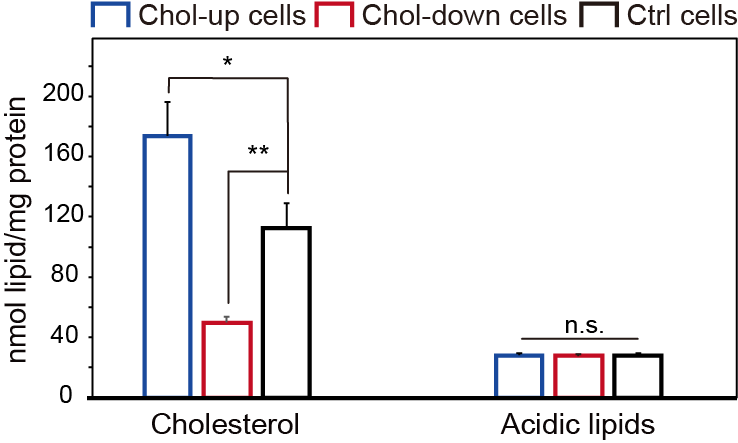
 Supplementary Fig. 7 | Amounts of cholesterol and acidic lipids in cholesterol-regulated cells. Lipid amounts were measured by MS-based lipidomic profiling. The acidic lipids include PS, PI, PG, PA. The detailed quantification data were shown in Supplementary Table 1. Error bars are standard deviations of three replicates. *, *p*-value < 0.05; **, *p*-value < 0.01; n.s. represents “not significant”. *p*-Values were analyzed by Student’s t-test.

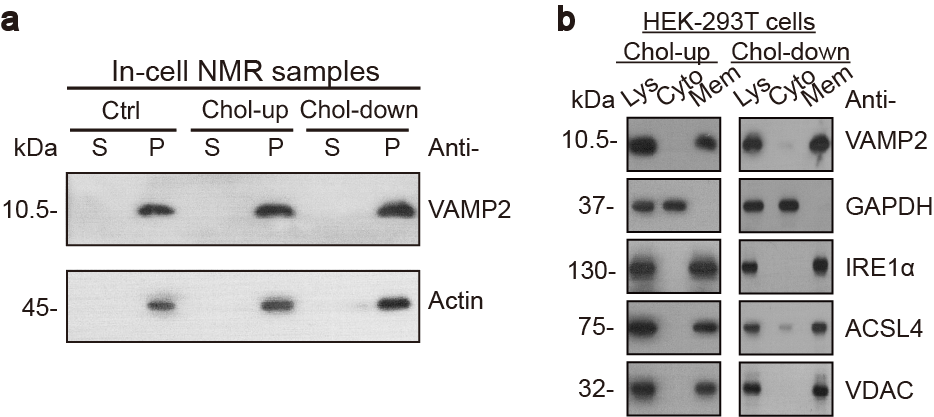
 Supplementary Fig. 8 | Amounts and localization of delivered VAMP2 in cholesterol-regulated cells. a, Comparisons of the leakage of VAMP2(1-96) in Chol-up, Chol-down and control HEK-293T cells by immunoblotting. The supernatant medium (S) and cell pellets (P) were collected after in-cell NMR experiments. b, Localization of delivered VAMP2(1-96) in Chol-up and Chol-down cells. Fractions of the total lysates (Lys), cytosol (Cyto) and membrane (Mem) were validated by immunoblotting with antibodies noted in Fig. 1D.

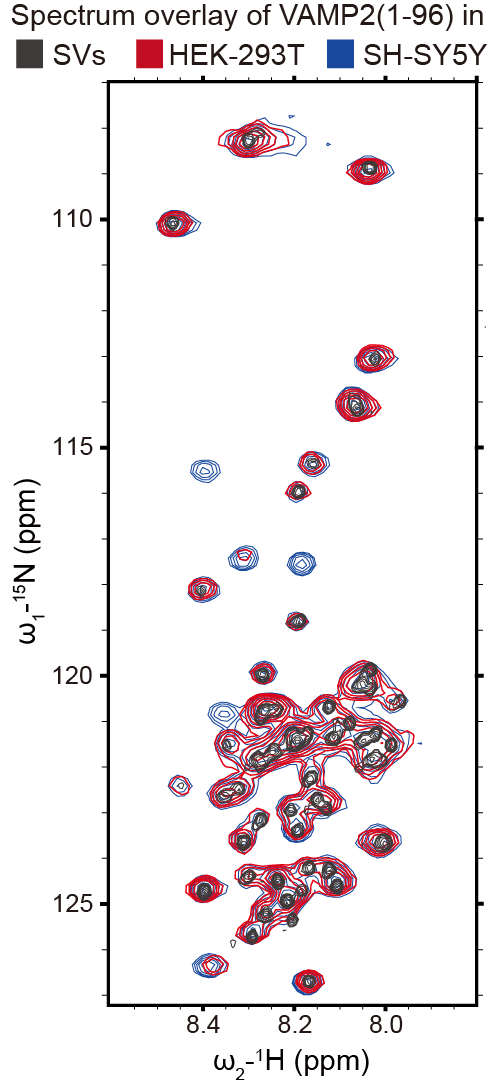
 Supplementary Fig. 9 | Overly of the 2D ^1^H-^15^N NMR spectra of VAMP2(1-96) with SVs (molar ratio is 700: 12), in HEK-293T cells and in SH-SY5Y cells.

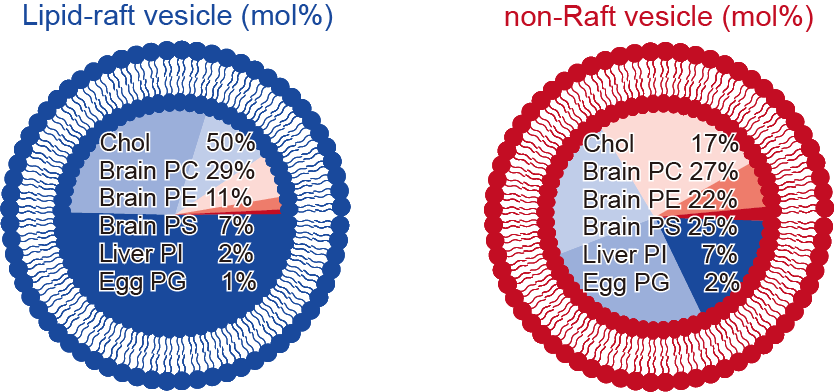
 Supplementary Fig. 10 | Lipid compositions of reconstituted lipid-raft- and non-raft-mimicking vesicles.

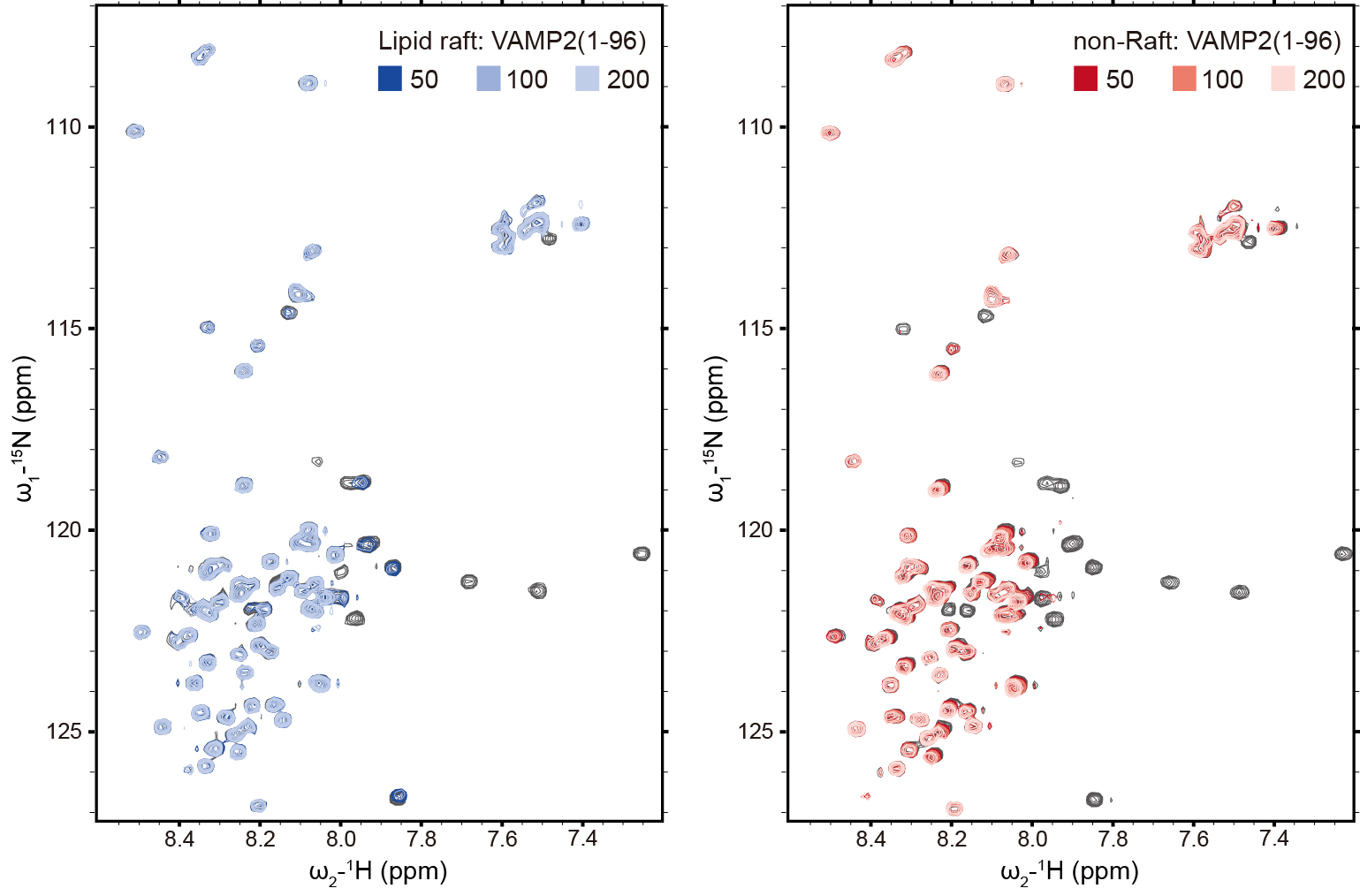
 Supplementary Fig. 11 | NMR titration of VAMP2 by lipid-raft- and non-raft-mimicking vesicles. 2D ^1^H-^15^N NMR spectra of VAMP2(1-96) are shown. The molar ratios of lipid to VAMP2(1-96) are indicated.

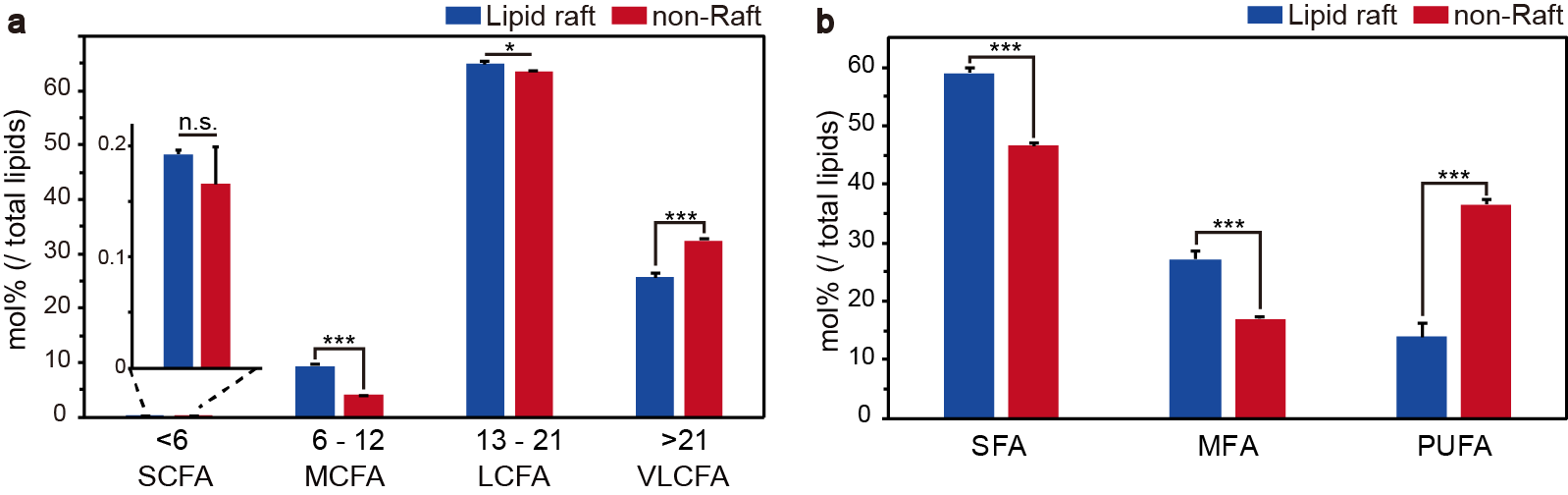
 Supplementary Fig. 12 | Comparisons of the acyl hydrocarbon tails of glycerophospholipids in lipid-raft and non-raft membranes. a, The acyl hydrocarbon tails were categorized by length into four classes (carbon numbers are indicated). SC: short-chain; MC: middle-chain; LC: long-chain; VLC: very long-chain; FA: fatty acid. b, The acyl hydrocarbon tails were categorized by saturation levels into three classes. S: saturated; M: monounsaturated; PU: polyunsaturated. Error bars are standard deviations from three biological replicates. *, p-value < 0.05; **, p-value < 0.01; ***, p-value < 0.001; n.s. represents “not significant”. p-Values were analyzed by Student’s t-test.

Supplementary Table 1. (separate Excel file) | Absolutely-quantitative identified lipids of cholesterol-level regulated cells. Chol-up: cholesterol-level up-regulated cells; Chol-down: cholesterol-level down-regulated cells; Ctrl: untreated control cells. Each cholesterol-level regulated cells were divided into three replicates for lipid quantification and normalized by the protein amounts. pPC: plasmenyl-phosphatidylcholine

Supplementary Table 2. (separate Excel file) | Absolutely-quantitative identified lipids in lipid-raft and non-raft membranes of SVs. Three separately isolated lipid-raft and non-raft samples were quantified by MS-based lipidomic profiling.

Supplementary Table 3. Abbreviation of lipid class |

PC	Phosphatidylcholine

pPC	plasmenyl-phosphatidylcholine

LPC	Lysophosphatidylcholine

aLPC	plasmanyl-lysophosphatidylcholine

PE	Phosphatidylethanolamine

pPE	plasmenyl-phosphatidylethanolamine

LPE	Lysophosphatidylethanolamine

pLPE	plasmenyl-lysophosphatidylethanolamine

PS	Phosphatidylserine

LPS	Lysophosphatidylserine

PG	Phosphatidylglycerol

LPG	lysophosphatidylglycerol

PI	Phosphatidylinositol

LPI	Lysophosphatidylinositol

PA	Phosphatidic acid

LPA	Lysophosphatidate

Cer	Ceramide

SM	Sphingomyelines

HexCer Hexosylceramide

DG	Diacylglycerols

TG	Triacylglycerols

Cho	Cholesterol
